# Maturation and persistence of the anti-SARS-CoV-2 memory B cell response

**DOI:** 10.1101/2020.11.17.385252

**Authors:** Aurélien Sokal, Pascal Chappert, Anais Roeser, Giovanna Barba-Spaeth, Slim Fourati, Imane Azzaoui, Alexis Vandenberghe, Ignacio Fernandez, Magali Bouvier-Alias, Etienne Crickx, Asma Beldi Ferchiou, Sophie Hue, Laetitia Languille, Samia Baloul, France Noizat-Pirenne, Marine Luka, Jérôme Megret, Mickaël Ménager, Jean-Michel Pawlotsky, Simon Fillatreau, Felix A Rey, Jean-Claude Weill, Claude-Agnès Reynaud, Matthieu Mahévas

**Affiliations:** Institut Necker Enfants Malades (INEM), INSERM U1151/CNRS UMS 8253, Université de Paris, Paris, France; Service de Médecine Interne, Centre Hospitalier Universitaire Henri-Mondor, Assistance Publique-Hôpitaux de Paris (AP-HP), Université Paris-Est Créteil (UPEC), Créteil, France; Inovarion, Paris, France; Institut Pasteur, Unité de Virologie Structurale, Paris, France; Département de Virologie, Bactériologie, Hygiène et Mycologie-Parasitologie, Centre Hospitalier Universitaire Henri-Mondor, Assistance Publique-Hôpitaux de Paris (AP-HP), Créteil, France; INSERM U955, équipe 18. Institut Mondor de Recherche Biomédicale (IMRB), Université Paris-Est Créteil (UPEC), Créteil, France; INSERM U955, équipe 2. Institut Mondor de Recherche Biomédicale (IMRB), Université Paris-Est Créteil (UPEC), Créteil, France; Département Immunologie-Hématologie, Centre Hospitalier Universitaire Henri-Mondor, Assistance Publique-Hôpitaux de Paris (AP-HP), Université Paris-Est Créteil (UPEC), 94000, Créteil, France; INSERM U955, équipe immunorégulation et biothérapie (I-BIOT) Institut Mondor de Recherche Biomédicale (IMRB), Université Paris-Est Créteil (UPEC), Créteil, France; Institut de Recherche Vaccinale (VRI), Université Paris-Est Créteil (UPEC), Faculté de Médecine, Créteil, France; INSERM U955, équipe 16. Institut Mondor de Recherche Biomédicale (IMRB), Université Paris-Est Créteil (UPEC), Créteil, France; Département de Santé Publique, Unité de Recherche Clinique (URC), CEpiA (Clinical Epidemiology and Ageing), EA 7376-Institut Mondor de Recherche Biomédicale (IMRB), Centre Hospitalier Universitaire Henri-Mondor, Assistance Publique-Hôpitaux de Paris (AP-HP), Université Paris-Est Créteil (UPEC), Créteil, France; Etablissement Français du Sang, INSERM U955, Université Paris-Est Créteil (UPEC), Créteil, France; Réponses inflammatoires et réseaux transcriptomiques dans les maladies, Institut Imagine, INSERM UMR1163, ATIP-Avenir Team, Université de Paris, Paris, France; Labtech Single-cell@Imagine, Institut Imagine, INSERM UMR 1163, Paris, France; Plateforme de Cytométrie en Flux, Structure Fédérative de Recherche Necker, INSERM US24-CNRS UMS3633, Paris, France

## Abstract

Memory B cells play a fundamental role in host defenses against viruses, but to date, their role have been relatively unsettled in the context of SARS-CoV-2. We report here a longitudinal single-cell and repertoire profiling of the B cell response up to 6 months in mild and severe COVID-19 patients. Distinct SARS-CoV-2 Spike-specific activated B cell clones fueled an early antibody-secreting cell burst as well as a durable synchronous germinal center response. While highly mutated memory B cells, including preexisting cross-reactive seasonal Betacoronavirus-specific clones, were recruited early in the response, neutralizing SARS-CoV-2 RBD-specific clones accumulated with time and largely contributed to the late remarkably stable memory B-cell pool. Highlighting germinal center maturation, these cells displayed clear accumulation of somatic mutations in their variable region genes over time. Overall, these findings demonstrate that an antigen-driven activation persisted and matured up to 6 months after SARS-CoV-2 infection and may provide long-term protection.

## Introduction

The new emerging coronavirus, SARS-CoV-2, has infected 55 million people and killed over 1 million individuals worldwide since the beginning of the pandemic. Understanding the mechanisms underlying the establishment of protective immune memory in recovering individuals is a major concern for public health and for anticipating vaccination outcomes.

In response to viral infection, virus-specific T and B cells are activated, expand and differentiate into effector cells (Wherry and Ahmed, 2004). At the early phase of COVID-19 infection, most infected individuals generate antibodies targeting the viral nucleocapsid (N) and the spike (S) proteins of SARS-CoV-2, including spike receptor binding domain (RBD) specific antibodies with strong neutralizing potential (Wajnberg et al., 2020). COVID-19 patients also develop potent early CD8^+^ and CD4^+^ T cell specific responses (Braun et al., 2020; Ferretti et al., 2020; Grifoni et al., 2020, 2020; Le Bert et al., 2020; Meckiff et al., 2020; Peng et al., 2020; Sekine et al., 2020; Swadling and Maini, 2020)

Studies have documented that B cells from patients with severe COVID-19 harbored low mutations frequencies in their heavy-chain variable region (VH) genes, notably those producing anti-RBD antibodies (Nielsen et al., 2020). This suggested an active early extrafollicular response in which naïve unmutated B cells first engage in cognate interactions with T cells and become fully activated to divide and differentiate into plasma cells (Jenks et al., 2019). Along with limited somatic hypermutation (SHM) and selection, cells derived from extrafollicular responses are usually considered as short-lived (Mac Lennan et al., 2003).

Long-term humoral immunity following infection relies on two types of cells derived from germinal center responses: long-lived plasma (LLPC), which continuously secrete antibodies (Slifka et al., 1998) and memory B cells (MBCs), which can expand and differentiate into antibody-secreting cells (ASC) upon a new antigenic challenge. MBCs can also adapt to cope with antigenic changes linked to virus mutations, as exemplify by the coevolution of neutralizing antibodies against mutating epitopes during HIV infection (Escolano et al., 2016; Liao et al., 2013; Wu et al., 2011). Assessing whether a MBC response, with the ability to rapidly produce SARS-CoV-2 neutralizing antibodies upon a new infectious challenge months or years after initial infection, is indeed established in convalescent COVID-19 patients is thus of immediate relevance for ongoing modeling of herd immunity and the formulation of vaccine strategies.

Serological studies have so far reported contradictory results on the persistence of humoral immunity in asymptomatic, mild and severe patients (Iyer et al., 2020; Long et al., 2020; Luchsinger et al., 2020; Pierce et al., 2020; Ripperger et al., 2020; Wajnberg et al., 2020; Weisberg et al., 2020). Furthermore, although memory B cells have been observed up to 6 months after infection in an increasing number of recent publication (Chen et al., 2020; Gaebler et al., 2020; Nguyen-Contant et al., 2020; Rodda et al., 2020; Vaisman-Mentesh et al., 2020), a recent study suggested that severe SARS-CoV-2 infection may blunt the germinal center (GC) response and subsequently compromise the generation of long-lived, affinity-matured memory B cells (Kaneko et al., 2020). In this context, analysis of the longevity and functionality of the anti SARS-CoV-2 memory B cell response becomes a major issue.

In the present study, we present a longitudinal deep profiling of the anti-SARS-CoV-2 memory B cell response in two parallel cohorts of patients with severe (S-CoV) and mild (M-CoV) COVID-19. We combined single cell transcriptomics, single cell culture and IgH VDJ sequencing to track and characterize the cellular and molecular phenotype and clonal evolution of spike-specific MBCs clones from early time points after SARS-CoV-2 infection up to 6 months after the initial symptoms. Our results provide new insights on the origin, magnitude and stability of the anti-SARS-CoV-2 MBC response and revealed in most patients a robust acquisition of GC-derived humoral immunity.

### Investigating anti-SARS-CoV-2 humoral response in mild and severe COVID-19 patients

To characterize the longitudinal evolution of the humoral response against SARS-CoV-2, we set-up a prospective cohort of patients with severe (S-CoV, n=21) and mild forms of COVID-19 (M-CoV, n=18) (MEMO-COV, NCT04402892, see **Methods and Table S1**). Initial samples were collected from patients in median 18.2 days and 35.5 days after disease onset for S-CoV and M-CoV respectively (M0). Two additional blood samples were collected at 3 (M3), and 6 months (M6), at a stage when all patients had fully recovered. All sera were first examined for SARS-CoV-2-specific IgG antibodies against the immunodominant nucleocapsid (N) and spike (S) proteins using commercial ELISA. As previously observed in other studies (Ripperger et al., 2020), N-specific IgG antibodies showed a rapid decrease in both cohorts (**Figure 1A)**. This decrease was particularly pronounced in M-CoV patients, among which 9 out of 18 had titers below the threshold of positivity at 6 months (**Figure 1A, Table S1**). In contrast, S-specific IgG antibodies appeared quite stable with time and only two M-CoV patients showed anti-S antibodies titers below detection threshold at the time of our last sampling (**Figures 1B and 1C**). Levels of S-specific IgG antibodies at 3 and 6 months, however, were significantly higher in patients who had developed a more severe form of the disease (**Figures 1B-1C**). Elevated titers of anti-S IgG strongly correlated with enhanced neutralization potential of patients’ sera in vitro (**Figure 1D, Figures S1A and S1B**), confirming the spike trimetric glycoprotein as a primary target of the protective immune response in patients.

**Figure 1.**
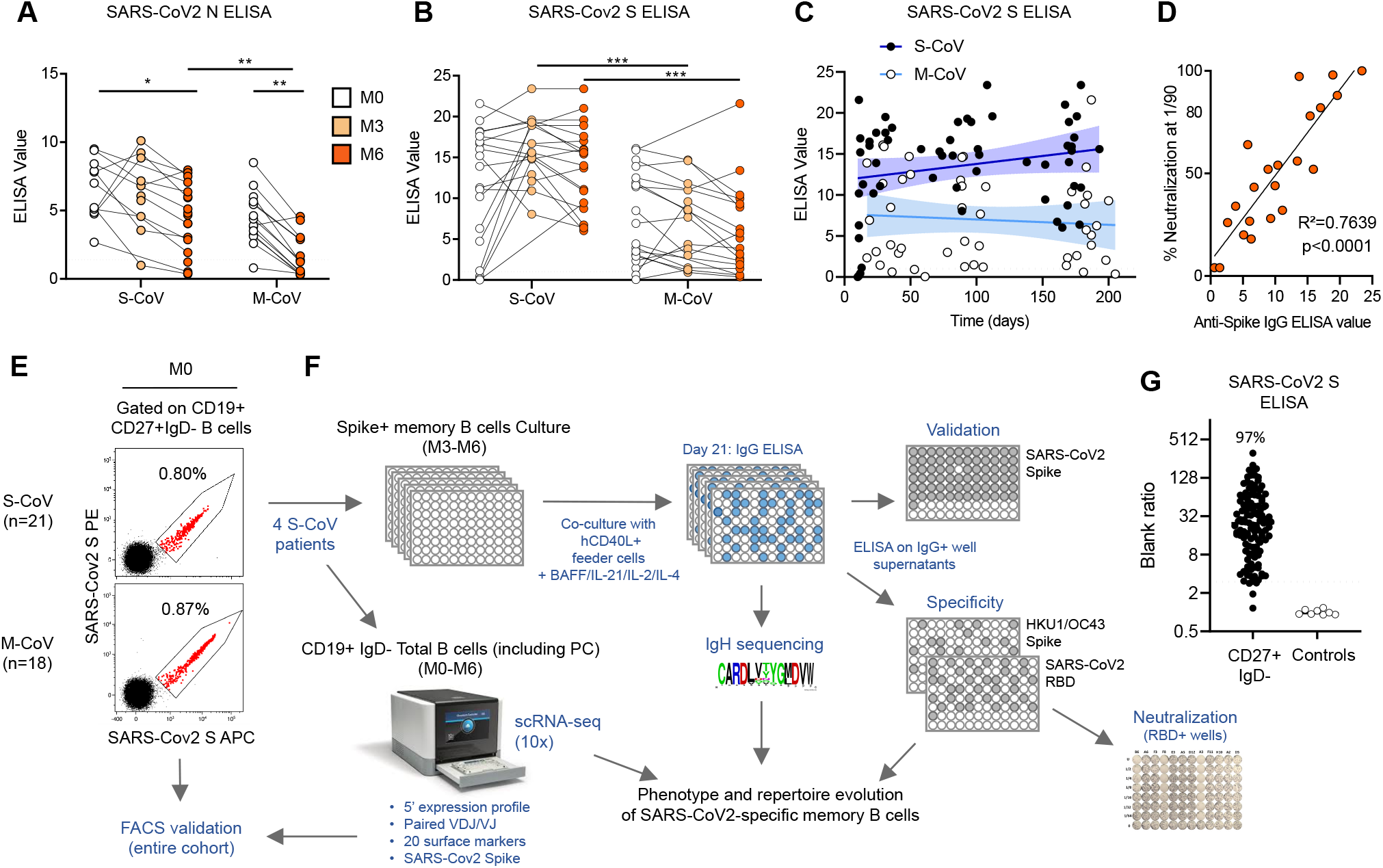
Longitudinal characterization of the humoral response against SARS-CoV2 in severe and mild COVID-19 patients. **(A-B)** Anti-SARS-CoV2 nucleocapsid (N) **(A)** and spike (S) **(B)** serum IgG levels were measured by ELISA in 21 severe COVID-19 (S-CoV) and 18 mild-COVID-19 (M-CoV) at M0 (white), M3 (light orange) and M6 (dark orange). The dashed line indicates the positivity threshold provided by the manufacturer. **(C)** Evolution of anti-SARS-CoV2 S serum IgG levels over time post-onset of COVID-19 symptoms in S-CoV (black dots, dark blue line) and M-CoV (white dots, light blue line) patients. Continuous lines indicate linear regression, colored area between dashed-lines indicates error bands (R²=0.049 for S-CoV, ns and 0.0061 for M-CoV, ns, Pearson correlation). **(D)** Correlation between sera anti-SARS-CoV2 S IgG levels and in vitro neutralization potential (% neutralization achieved at a 1/90 dilution) at M6 (n=10 S-CoV and 11 M-CoV patients). The line represents a simple linear regression (R² and p-value with Pearson correlation are shown) **(E)** Representative FACS plot of His-tagged SARS-CoV2 S staining in gated live CD19^+^CD38^int/-^CD27^+^IgD^−^ B cells at M0 in two representative S-CoV (upper plot) and M-CoV patients (lower plot). **(F)** Overall study design. **(G)** Representative results of an anti-SARS-CoV2 S IgG ELISA on supernatants from sorted SARS-CoV2 S-specific MBCs (dark dots) and non spike-specific MBCs (white dots), as validation of FACS S-Cov staining. Lines indicate median value. Dashed line indicates the positivity threshold (≥3 x blank). Anova and two-tailed Mann-Whitney tests were performed (**P < 0.01, *P < 0.05).

To further track the SARS-CoV-2-specific memory B cell response, we next implemented two complementary in-depth approaches on four S-CoV patients’ samples. First, we generated His-tagged trimeric SARS-CoV-2 spike ectodomain and used successive staining with His-tagged S and two fluorescently labeled anti-His antibodies to stain and perform high purity single-cell sorting of SARS-CoV-2 S-specific B cells (**Figure 1E**). Single B cells were further cultured in an optimized in vitro culture assay (McCarthy et al., 2018) (**Figure 1F**). After 21 days of culture, RNA was extracted from single-cell cultures to determine their immunoglobulin heavy chain (IgH) sequence. Culture supernatants with IgG concentration over 1μg/ml were additionally tested by ELISA to validate the cell specificity towards SARS-CoV-2 S (over 95 % purity in this assay (**Figure 1G**)), investigate its recognition of SARS-CoV-2-S receptor binding domain (RBD) and neutralization potential, and analyze potential cross-reactivity to seasonal coronaviruses (HCoV-HKU1/HCoV-OC43). In parallel, to get a broader view of the overall B cell response towards SARS-CoV-2 at both early (M0) and late time points (M6), we sorted CD19^+^IgD^-^ B cells from the same patients (**Figure S1C)** and performed scRNA-seq using the 10X Genomics technology. This allowed us to couple both 5′ single-cell RNA sequencing, V(D)J profiling and Feature Barcoding of 20 known B cell surface markers, and we developed as well anti-His barcoded antibodies to reveal His-tagged SARS-CoV-2 S-specific B cells.

### Spike-responding B cells harbor phenotypically and transcriptionally an activated phenotype and a time-dependent maturation toward the memory B cell lineage

Unsupervised clustering analysis of our scRNA-seq dataset revealed that CD19^+^IgD^-^ B cells could be divided in 5 major clusters according to their gene expression profile (**Figure 2A-B and S2A-B**), a clustering which correlated nicely with the expression of barcoded surface markers included in our dataset (**Figure 2C-D**). Two of these clusters corresponded to antibody-secreting cells (ASCs), with both a short-lived plasmablasts cluster (PB), enriched for expression of cell proliferation-related genes, and non-dividing plasma cells (PC). Both PB/PC populations were strongly reduced by 6 months, confirming the resolution with time of the primary extrafollicular response in these severe COVID-19 patients (**Figures 2B**). The remaining B cells were separated in 3 populations: a mixture of naïve/transitional B cells, a resting memory B cell population (MBCs) and a CD95^+^ activated cluster. This activated cluster could be further subdivided in 3 distinct populations: CD21^low^CD27^+^CD38^+^CD71^+^ activated B cells (ABC), CD21^low^CD27^low^CD38^−^CD71^low^CD11c^+^FcRL5^+^ cells, likely corresponding to atypical memory and/or double-negative 2 population (DN2) (Sanz et al., 2019) and CD21^+^CD27^int/+^CD38^−^CD71^low^CD95^+^ cells corresponding to a cluster with intermediate characteristics between ABC and memory B cells (**Figures 2C-D and S2A-B**). The contraction of the primary response, clearly observed for ASCs, could also be seen for ABCs, with a parallel increase in resting MBCs. All other clusters remained stable.

**Figure 2.**
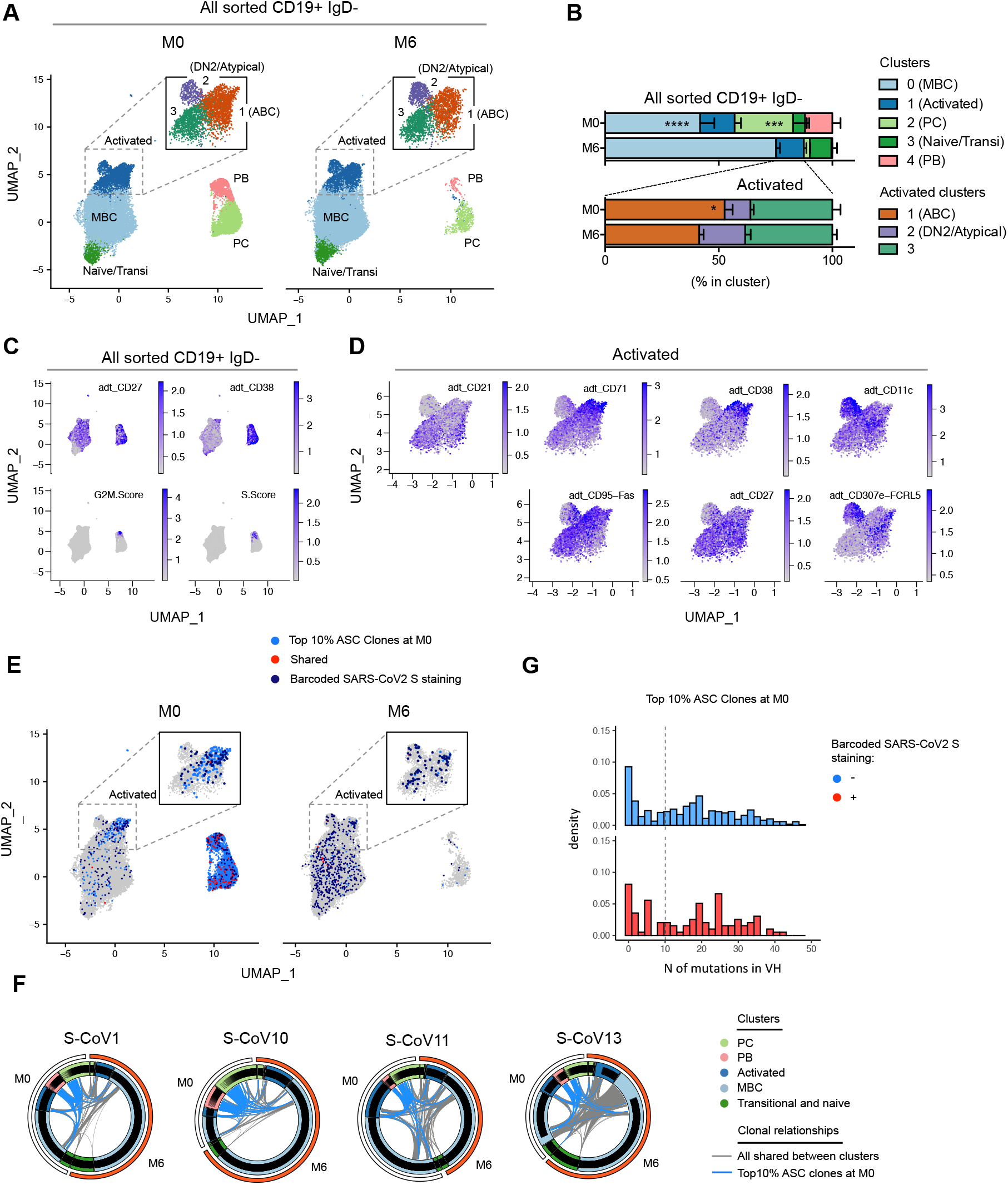
Characterisation of the B response against SARS-CoV-2 in acute (M0) and convalescent (M6) severe COVID-19 patients. **(A)** UMAP and clustering of 41083 B cells analysed by scRNA-seq from 4 S-CoV patients at M0 (left panel) and M6 (right panel) (see Table S2). Upper square in both panels shows the results of increased clustering resolution for the “Activated” cell cluster. **(B)** Relative cluster distribution at M0 and M6 for all sorted cells (Top panel) and cells falling in the “Activated” cluster at initial clustering resolution. Bar indicates mean with SEM. **(C)** Feature Plots showing scaled normalized counts for CD27 and CD38 barcoded antibodies as well as S score and G2/M signature scores in all cells. **(D)** Feature plots showing scaled normalized counts for CD21, CD71, CD38, CD11c, CD95, CD27 and CD307e (FcRL5) barcoded antibodies in cells from the “Activated” cluster. **(E)** UMAP of all cells at M0 or M6, with cells belonging to one of the 10 percent most expanded antibody secreting cell (ASC) clones highlighted (light blue). Positive cells for barcoded-SARS-CoV2-S staining are also highlighted (red when members of one of the 10% most expanded ASC clones at M0, dark blue otherwise). **(F)** Circus plot showing clonal relationships between cells from different UMAP clusters at M0 and M6. Blue lines indicate clones belonging to the top 10% ASC clones at M0 and grey line all other shared clones. **(G)** Histograms showing the number of mutations in V genes for cells belonging to one of the 10% most expanded ASC clones at M0 according to their barcoded-SARS-CoV2-S staining.

To gain insight into the specificity of this B cell response and the clonal relationship between the different B cell clusters, we first used the initial burst of ASCs seen at M0 as a proxy for the overall SARS-CoV-2-specific B cell response. Investigating the top 10% ASC clones at M0 for each patient in our dataset revealed a strong link between the ABC cluster and the ASC population (**Figures 2E-F and S2C-D**). The bimodal distribution of somatic hypermutation (SHM) seen in ASCs further suggests the existence of a strong extra-follicular reaction recruiting newly engaged naïve and memory B cells, which fuels the primary response towards SARS-CoV-2 (**Figure 2G**). In line with the resolution of the immune response, most of these clones were no longer present in ASCs six months after SARS-CoV-2 infection, with only few cells, clonally-related to the original ASC burst, still detectable in residual PC, while they were increased in the resting MBC compartment by that time (**Figures 2E and 2F, S2C and S2D)**. Parallel analysis of barcoded-spike reads in our dataset allowed us to identify clear populations of Spike-specific ABCs and ASCs at M0, mirroring - yet showing limited overlapping with - the bulk of the ASC primary response. By 6 months, spike-specific cells mainly localized in the resting MBC cluster and showed significant expansion with still very limited relationship with the original ASC response, suggesting two distinct, albeit synchronous responses in COVID-19 patients.

### Spike-specific memory B cells mature from ABCs and accumulate up to 6 months pos-infection in COVID-19 patients

To further analyze the dynamics of the overall and SARS-CoV-2 S-specific B cell response at the level of our entire cohort, we next performed multi-parametric FACS analysis using a flow panel which included 7 surface markers identified in our scRNA-seq dataset as sufficient to clearly delineate all major B cell and ASC subsets involved in the initial response (CD19, CD21, CD27, CD38, CD71, CD11c, IgD, as well as His-tagged trimeric S protein). Unsupervised analysis of CD19^+^IgD^−^ switched B cell populations (**Figures S3A-C**), concatenated from 83 samples collected at M0, M3 and M6 from patients in our cohort, confirmed our initial observation that Spike-specific B cells were enriched in a cluster of CD27^+^CD38^int/+^CD71^+^ ABCs and in ASCs at M0 (**Figures 3 A-C**). The enrichment in ASCs was most prominent in severe COVID-19 patients and is in line with the strong ASC burst seen in these patients (**Figures S3 A-C).**Spike-specific ASCs were marginally detectable at 6 months, at which time point Spike-specific B cells mostly resided in the CD21^+^CD27^+^CD38^−^ CD71^int/-^ resting memory B cell compartment in both severe and mild COVID-19 convalescent patients. Of note, some spike-specific cells were still found in the CD71^+^ ABC cluster at M6 in both groups of patients but were much less frequent in the DN2/atypical compartment **(Figure 3C)**.

**Figure 3.**
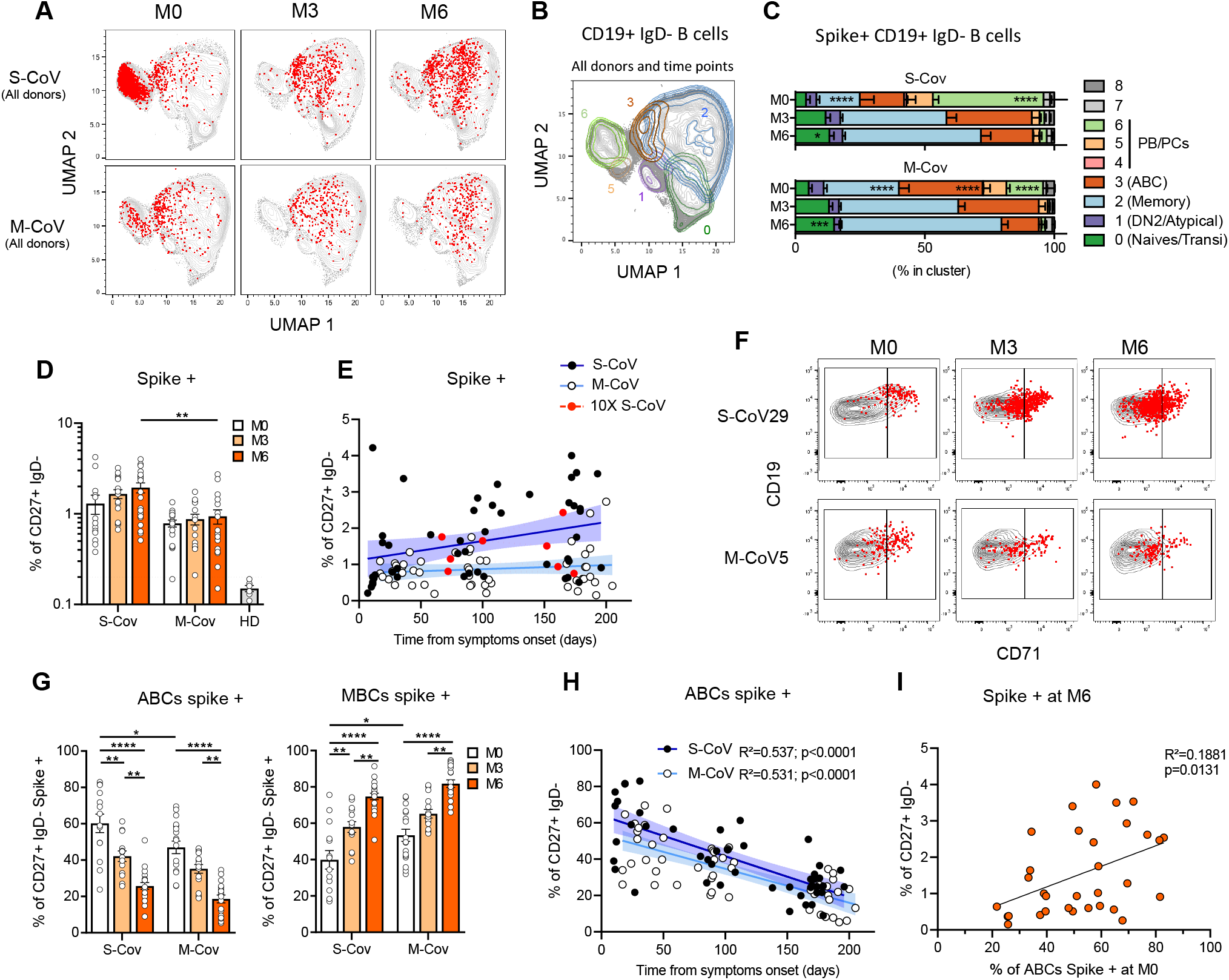
Phenotypic evolution of the SARS-Cov2 S-specific B cell response up to six months post infection in mild and severe COVID-19 patients. **(A)** UMAP projections of concatenated COVID-19 multi-parametric FACS analysis of CD19^+^IgD^−^ cells from all S-CoV (n=15) and M-CoV (n=16) patients analyzed over time in our cohort (Table S1). His-tagged labeled SARS-CoV2 S-specific cells are overlaid as red dots. **(B-C)** Unsupervised clustering (Flow SOM) was performed on the concatenated FACS dataset based on IgD, CD71, CD27, CD38, CD11c, CD19, CD21 fluorescence intensity. Main defined clusters (>5% of total CD19^+^IgD^−^ B cells) are shown as overlaid contour plots on the global UMAP representation **(B)**. Cluster distribution of SARS-CoV2 S-specific cells in identified clusters, at indicated time point, is further displayed as bar plots **(C)**. Bars indicate mean with SEM. **(D)** Proportion of SARS-CoV2 S-specific CD19^+^IgD^−^CD27^+^CD38^−^ MBCs at M0, M3 and M6 in S-CoV and M-CoV patients, compared to 6 pre-pandemic healthy donor controls (HD). Each dot represents one patient, bars indicate mean with SEM. Dashed line indicated the mean + 2 standard deviations of the detected frequency of SARS-CoV2 S-specific MBCs in HD. **(E)** Proportion of spike-specific MBCs over time post onset of COVD-19 symptoms in S-CoV (black dots, dark blue line) and M-CoV (white dots, light blue line) patients. Lines indicates linear regression (r²=0.051, p-value=0.02 for S-CoV, ns, and r²=0.018 for M-CoV, ns. Pearson correlation). Colored area between dashed-lines indicates error bands. Red dots and lines indicate the 4 S-CoV patients analyzed as part of our scRNA-seq dataset. **(F)** Representative dot plot showing CD71 and CD19 expression in IgD^−^CD19^+^CD38^−^ B cell at indicated time points from two representative S-CoV and M-CoV patients. SARS-CoV2 S-specific cells are overlaid as red dots. Gating strategy for ABCs and classical MBCs according to CD71 expression is further displayed. **(G)** Proportion of SARS-CoV2 S-specific displaying an ABC (CD19^+^CD27^+^IgD^−^CD71^+^) (left) or MBC (CD19^+^CD27^+^IgD^−^CD71^low/int^) (right) phenotype at indicated time points. Bars indicate mean with SEM **(H)** Proportion of activated MBCs over time in S-CoV (black dots, dark blue line) and M-CoV (white dots, light blue line) patients. Lines represents the linear regression. Colored area between dashed-lines indicates error bands. R² and p-value with Pearson correlation. **(I)** Correlation between the frequency of SARS-CoV2 S-specific MBCs at M6 and of ABCs at M0 in all M-CoV and S-CoV patients. Linear regression with Pearson correlation analysis. Anova and two-tailed Mann-Whitney tests were performed (**** P < 0.0001, **P < 0.01, *P < 0.05).

Based on these results, and using a more traditional gating strategy, we next focused our analysis on the switched CD27^+^IgD^−^CD38^int/-^ memory B cell compartment, which includes both ABCs and resting memory B cells (**Figure S1C**). In contrast to the rapid disappearance of spike-specific ASCs, both the percentage and absolute number of spike-specific CD27^+^IgD^−^ B cells appeared stable up to 6 months in the vast majority of patients in our cohort, and even continuously increased up to that time point in a subset of the convalescent S-CoV patients (**Figures 3D-E and S3D and S3D**). Of note, and as previously mentioned, the double-negative population contained few spike-specific cells (**Figures S3F and S3G**). Convalescent S-CoV patients showed significantly higher frequencies of spike-specific MBCs at the 6-month time point of our study. Most M-CoV patients, however, still harbored a sizeable population of spike-specific memory B cells at 6 months post-infection (mean 0.94% ±0.17 of MBCs) and only one (M-CoV24) out of the 16 M-CoV patients analyzed at 6 months in our study showed a frequency of spike-specific switched CD27^+^ memory B cells below that of pre-pandemic healthy donors. Even more striking is that both M-CoV patients whose serum levels of S-specific IgG had drop below detectable levels by 6 months (correlating with an absence of in vitro neutralizating potential) still harboured a clear population of spike-specific MBCs at that time point (0.41 and 0.60% of MBCs, respectively). Altogether, these results demonstrate the induction of a robust and stable spike-specific MBC population in both mild and severe COVID-19 patients. Aside from severity, none of the clinical parameters or treatment that we could monitor, notably corticosteroids, appeared to have any clear influence on the long-term establishment of these spike-specific MBCs (**Figures S3J-N).**

To ensure that such stable levels of spike-specific CD27^+^ switched-memory B cells did not simply reflect a sustained immune response in these convalescent COVID-19 patients, we further used CD19 and CD71 to separate ABCs (CD19^high^CD71^+^) and resting MBCs (CD19^+^CD71^low^) among spike-specific cells over time, as previously described (Ellebedy et al., 2016). Confirming our initial observation in the 4 S-CoV patients included in the scRNAseq dataset (**Figure 2E**) and the clear enrichment of spike-specific B cells in cluster 3 of our unsupervised FACS analysis (**Figures 3A-C**), the majority of spike-specific memory B cells displayed a CD19^high^CD71^+^ ABC phenotype in the days following infection (**Figure 3F).**In both S-CoV and M-CoV, the proportion of spike-specific ABCs steadily decreased over time, along with an increase of spike-specific classical, resting MBCs (**Figures 3 G and 3H)**. Interestingly, ABCs were still detectable in both cohorts at 6 months (approximately 20% in both cohorts), suggesting continued antigen-driven activation, as has been described after Ebola infection (Davis et al., 2019a). In S-CoV patients, a small fraction of these early responding MBCs expressed the transcription factor T-bet, but such expression appeared only transient (**Figure S3H**). More interestingly, we noted a significantly higher frequency of ABCs among spike-specific MBCs in S-CoV as compared to M-CoV patients, as well as large inter-patients variability for this parameter in both group (**Figure 3 G)**. This heterogeneity could not be simply explained by inter-patient variations in the time of initial sampling. However, the proportion of spike-specific ABCs at M0 correlated significantly with the proportion of spike-specific MBCs at 6 months (**Figure 3I**). Overall, these results suggest a potential link between early activated B cells and memory B cells established at later time points in the response. They also highlight the presence of CD19^high^CD71^+^ early ABCs as a potential marker to predict the magnitude of classical, possibly long-term memory against SARS-CoV-2.

### Acquisition of somatic hypermutations in RBD-specific IgVH genes of MBCs

In order to better understand the origin, fine specificity and developmental pathways involved in the generation of memory B cells against SARS-CoV-2, we next performed single cell sorting and culture of spike-specific memory B cells at 3 and 6 months post-infection in all 4 donors previously analyzed in the context of our scRNA-seq experiment (**Figure 1E**). A total of 2412 cells at M3 and 2423 cells M6 from the 4 donors were sorted (**Table S2**), and 488 and 1027 supernatants (respectively at M3 and M6) post single-cell culture were tested for specificity against SARS-CoV-2 S protein as well as Spike protein from two other related seasonal Betacoronavirus (HCoV-HKU1 and HCoV-OC43) (Huang et al., 2020), and against the RBD domain of the SARS-CoV-2 spike protein which is the main target of neutralizing antibodies in COVID-19 patients. We found that a sizeable proportion of SARS-CoV-2 spike-specific memory B cells also recognized the spike protein of H-CoV-HKU1 and H-CoV-OC43, with many of these cells recognizing both, as previously reported (Shrock et al., 2020, Wec et al., 2020). The proportion of HCoV-HKU1 and HCoV-OC43 specific MBCs clones, however, readily decreased between M3 and M6 (12.4%±4.9 to 4.8%±1.7 respectively), suggesting that these specificities, while actively participating in the early response, are not preferentially recruited in the later memory B cell compartment. In contrast, the proportion of SARS-CoV-2 RBD-specific MBCs clones clearly increased between the M3 and M6 time points in all donors (11.1%±2.1 and 26.3%±5.1 respectively). (**Figures 4A and 4B, Figure S4A and S4B**). SARS-CoV-2 RBD-specific clones were mostly sorted from resting CD71^−^ CD27^+^ switched memory B cells at these late time points (**Figures S4C and S4D**) and were detectable within the same proportion in M-CoV and S-Cov patients at 6 months (**Figure 4B**). Finally, only a minority of SARS-CoV-2 RBD-specific clones cross-reacted with HCoV-HKU1 and HCoV-OC43 spike protein, as anticipated by the low similarity of the RBD domain between these different viruses (**Figures S4A and S4B**). Overall, these results suggested an early and rapid recruitment of cross-reactive memory B cells through the extra-follicular pathway, in parallel with the establishment of a new immune response against epitopes unique to SARS-CoV-2, with a delayed and progressive output of memory B cells.

**Figure 4.**
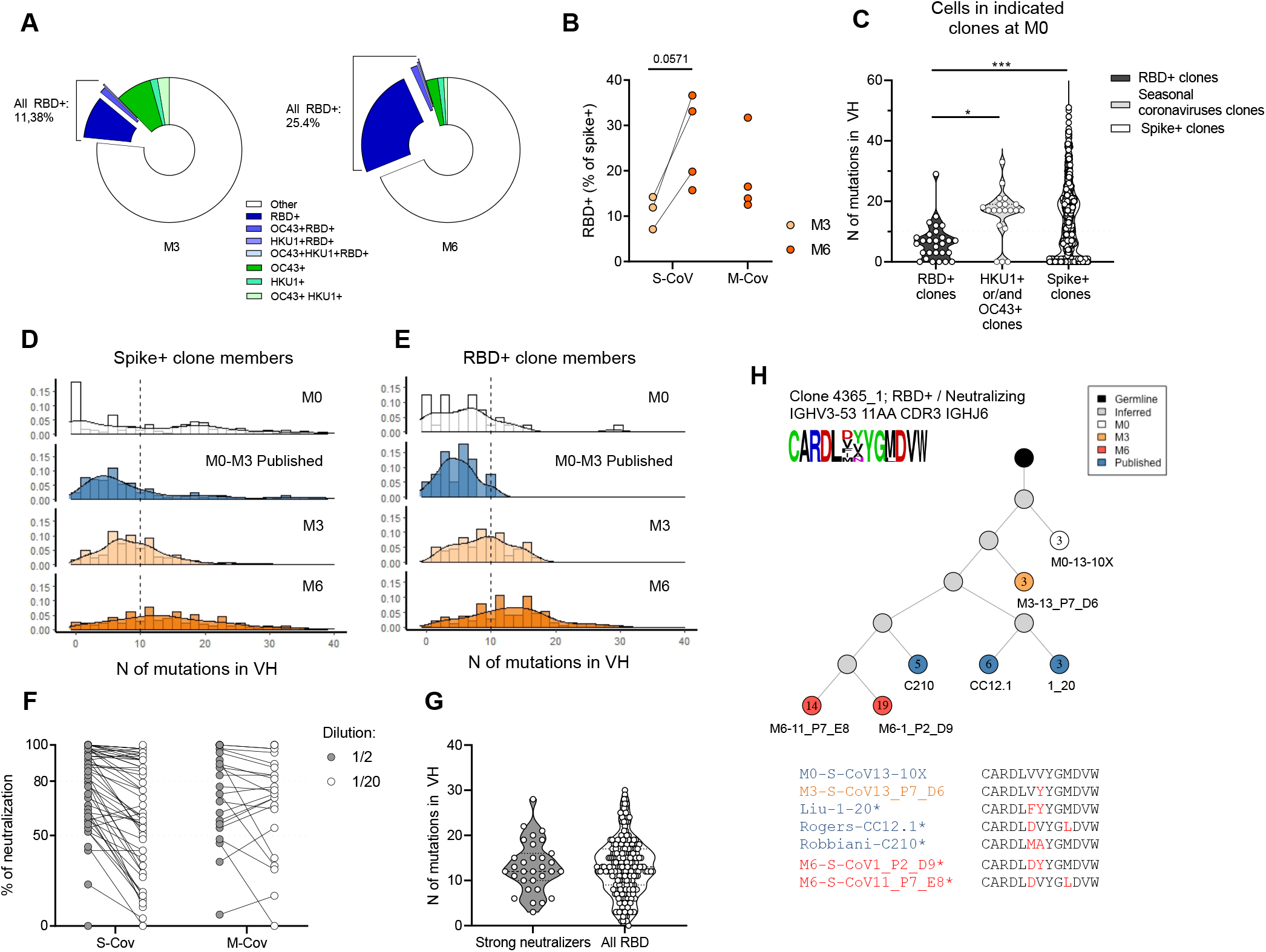
Maturation of the SARS-Cov2 S-specific repertoire up to six months post infection in mild and severe COVID-19 patients. **(A)** Pie charts showing the average percentage of RBD^+^ and cross-reactive specificity (RBD^+^OC43^+^, RBD^+^HKU1^+^OC43^+^, RBD^+^HKU1^+^, OC43^+^, HKU1^+^, OC43^+^HKU1^+^) among single-cell cultured spike-specific B cells as determined by ELISA. Three and 4 S-CoV patients, all previously included in our scRNA-seq dataset, were analyzed at M3 and M6, respectively. Average percentage of RBD specific cells among SARS-CoV2 S-specific sorted cells from indicated S-CoV and M-CoV patient at M3 and M6. **(C)** Violin plot showing the number of mutations in the Ig VH segment of cells in the original M0 10x scRNAseq VDJ dataset found to be in clonal relationship with M3 or M6 SARS-CoV2 S-specific sorted cells showing specificity against RBD (RBD^+^ clones), seasonal Beta-coronaviruses (HKU1-CoV or OC43-CoV) or against S only (Spike^+^ clones). **(D-E)** Histograms showing the relative distribution of mutation numbers in the Ig VH segment from all SARS-CoV2 S-**(D)** or RBD-**(E)** specific clone members (10x scRNA-seq dataset and single-cell heavy chain sequencing data) at M0, M3 and M6 as well as from sequences from the literature ((Brouwer et al., 2020; Kreer et al., 2020; Liu et al., 2020; Robbiani et al., 2020; Seydoux et al., 2020; Shi et al., 2020; Wec et al., 2020; Zost et al., 2020), mostly between M0 and M3). **(F)** Plot showing the percentage of neutralization from single cell culture supernatants of identified SARS-CoV2 RBD-specific B cells. Two dilutions (1/2 and 1/20) were assayed for each supernatant tested. Dashed line indicates 80% neutralization and 50% neutralization. **(H)** Violin plot representing the number of mutations in the Ig VH sequence in strong neutralizing antibodies at M6 (neutralization >80% at ½ dilution) *vs.* the number of mutations in the Ig VH sequence of all cells from anti-RBD clones at M6. **(H)** Evolutionary tree of an RBD-specific and neutralizing clone, built on sequences from 10X scRNA-seq, cell culture and literature. Each circle represents a unique sequence from that clone. Circle color indicates time-point of origin and the number inside indicates the calculated number of mutations from inferred germline. Grey indicates a theoretically inferred common progenitor. * indicate that the antibody associated with that sequence has been validated as neutralizing in vitro. CDR3 from all sequences in the tree are represented as a frequency plot logo (Top left) as well as below the tree, where each amino-acid in red indicates a change compared to the top listed CDR3.

To evaluate whether such memory B cell output corresponds to an ongoing GC response, we next evaluated the occurrence and evolution of somatic hypermutation (SHM) in spike-specific MBCs. Heavy chain VH sequencing from spike or SARS-CoV-2 RBD specific wells (**Table S2**) in our in vitro culture assay, at both M3 and M6 (**Figure S4E**), allowed us to identify clonal relationships with sequences obtained in our scRNA-seq dataset at both M0 and M6. Mutations in VH sequences of SARS-CoV-2 S-specific clones at M0 displayed a bimodal distribution with both near germline and highly mutated sequences. In line with an initial recruitment of cross-reactive memory B cells into the anti-SARS-CoV-2 early humoral response, HCoV-HKU1 and HCoV-OC43 specific clones contained sequences that were already highly mutated at M0 (**Figure 4C**). In contrast, SARS-CoV-2 RBD-specific clones displayed low SHM in VH sequences, as previously described in numerous studies during the early stages of this pandemic (Brouwer et al., 2020; Kreer et al., 2020; Liu et al., 2020; Robbiani et al., 2020; Seydoux et al., 2020; Shi et al., 2020; Wec et al., 2020; Zost et al., 2020). Both early SARS-CoV-2 S-specific and RBD-specific IgH sequences showed high level of convergence between donors in our dataset, but also with published SARS-CoV-2 spike- and RBD-specific antibody sequences (43/874 sequences in clonal relationship with cells from our dataset) (**Figure S4F**). Although near germline sequences were initially present in spike-specific clones, and even predominated in those with SARS-CoV-2 RBD specificity in both our M0 dataset and in the literature, longitudinal tracking of clonal relationships at 3 and 6 months revealed a progressive acquisition of SHM in VH sequences over time (**Figures 4D and 4E and S4G-H**). Most importantly, over 70% of SARS-CoV-2 RBD-specific culture supernatants at M6 showed intermediate to strong neutralizing potential against SARS-CoV-2 *in vitro* in both S-CoV and M-CoV patients (**Figure 4F**). Progressive accumulation of SHM in VH sequences was not associated with a loss of neutralization potential (**Figure 4H**). In contrast, a strong neutralizing capacity was identified along lineage trees including previously described near germline anti-SARS-CoV-2 RBD antibodies and the mutated versions that we identified at M6 in this work. (**Figures 4G and S4I**). Dynamics of such clones and emergence of highly mutated neutralizing sequences, along with the robustness of the memory response observed in S-CoV and M-CoV patients, demonstrate that MBCs bearing unique specificities and neutralizing potential against SARS-CoV-2 can expand and mature over time through an ongoing germinal center response.

## Discussion

The long-term stability of humoral immune memory following viral infection is provided by separate pools of protective long-lived plasma cells (LLPCs) and memory B cells that can last for a lifetime (Amanna et al., 2007). Upon re-infection, long-lived memory B cells generate a new wave of short-lived plasma cells and LLPC, while preserving the pool of memory B cells. In this study, we focused on SARS-CoV-2 infected patients with mild or severe disease and investigated the establishment of humoral memory and how this memory relates to the primary B cell response that emerges at the early stage of infection.

Experimental infections of volunteers with Betacoronaviruses responsible for common colds and longitudinal serological studies have suggested the absence of durable immunity against these seasonal infections (Sariol and Perlman, 2020). Furthermore, early reports have shown potentially defective GC formation in severe COVID-19 patients in intensive care unit (Kaneko et al., 2020) and predominantly near-germline anti-SARS-CoV-2 neutralizing antibodies (Ju et al., 2020; Nielsen et al., 2020; Robbiani et al., 2020). Such observations suggested a strong bias of the anti-SARS-CoV-2 immune response toward the extrafollicular pathway in these patients, which may preclude the formation of a long-term memory.

Our study demonstrate, in contrast, a remarkable stability of the overall spike-specific memory B cells population up to 6 months after infection, and even its expansion in severe patients, extending recent observations on memory persistence in COVID-19 (Gaebler et al., 2020). Tracking of spike-specific B cells provided a more complex picture than previously anticipated, characterized by two synchronous responses with distinct dynamics throughout the extra-follicular reaction. Indeed, near germline B cell clones recognizing the SARS-CoV-2 spike protein were mobilized, but also pre-existing highly mutated MBCs specific for the spike protein of other seasonal Betacoronaviruses.

The longitudinal VDJ sequencing of spike-specific MBCs revealed that neutralizing SARS-CoV-2 RBD-specific clones, including convergent antibody rearrangements across donors, accumulated with time and acquired somatic mutations in their variable region genes, the hallmark of a GC-dependent immune response. Such GC imprinting is of major importance because it usually predicts the generation of long-lived memory B cells (Mesin et al., 2016). Along this line, a recent report observed the presence of viral proteins in the intestine of patients several months after the initial diagnostic, which could sustain such B cell response (Gaebler et al., 2020).

Cross-reactivity against the spike domain of betacoronaviruses has been already documented in serological studies showing reactivity to conserved regions of the spike protein (Aydillo et al., 2020; Wang et al., 2020), by the presence of preexisting antibodies recognizing conserved epitopes in the S2 protein of SARS-CoV-2 in uninfected individuals (Ng et al., 2020) and cross-reactivity of sorted spike-specific B cells with seasonal coronavirus (Song et al., 2020; Wec et al., 2020). Such cross-reactivity also exists for T-cells (Braun et al., 2020; Grifoni et al., 2020; Le Bert et al., 2020), but its protective potential remains debated (Ng et al., 2020; Sermet et al., 2020). The proportion of cross-reactive clones, however, decreased from 3 to 6 months, suggesting that, after an initial activation, these clones were not positively selected in the memory pool.

As observed previously by others for antibody titers (Ripperger et al., 2020), the overall strength and stability of the memory B cell response was positively correlated with the initial severity of COVID-19-associated pathologies in convalescent patients. Interestingly, the memory response appeared also linked to the magnitude of the initial SARS-CoV-2 S-specific ABC response. Activated B cells in COVID-19 patients have been previously reported with various phenotypes (Juno et al., 2020; Mathew et al., 2020; Oliviero et al., 2020; Woodruff et al., 2020), including a double-negative CD11c-positive population (DN2). One study has also suggested the presence of SARS-CoV-2 RBD-specific DN2 cells as a correlate of critical illness (Kaneko et al., 2020). We found here, that most spike-responding B cells harbored an activated phenotype that notably differed from DN2 but was similar to the one described after Ebola or Influenza virus infection (Ellebedy et al., 2016). These SARS-CoV-2 spike-specific ABCs were in clonal relationship with expanded cells from the initial burst of antibody-secreting cells, but also with resting memory B cells that persisted at 6 months. Given the limited clonal overlap between the early ASC burst and the later mutated memory B cell response, it is tempting to suggest that a distinct part of ABCs is dedicated to fueling the GC response in these patients, while another part contribute to the ASC burst. Of note, the ABC population remains detectable up to 6 months post infection in convalescent patients, with some, albeit few spike-specific cells, suggesting, as described for Ebola (Davis et al., 2019), an antigen-driven activation that persisted for months after infection.

Collectively, taking into account this report and previous studies, the COVID-19 infection seems in most non-critical cases to induce both an immediate protective antibody response together with the ongoing maturation of memory B cells, which should give rise to neutralizing antibody-secreting cells upon reinfection. While this overall picture is positive, notably in the context of vaccine campaigns to come, the long-term duration of the protective memory B cell response remains an open question.

## Acknowledgments

We thank Garnett Kelsoe for providing us with the human cell culture system, together with invaluable advices. We thank the Prince of Monaco, and the Government Council for free donation. We also thank Sebastien Storck and Sandra Weller for their advices and support; we thank the physicians, Constance Guillaud, Fréderic Schlemmer, Elena Fois, Henri Guillet, Marc Michel, Bertrand Godeau, whose patients were included in this study.

## Funding

This work was supported by an ANR Grant (MEMO-COV-2 - FRM). AP-HP was the promotor and the sponsor was Assistance Publique – Hôpitaux de Paris (Département de la Recherche Clinique et du Développement). AS was supported by a Poste d’Accueil from INSERM, AR by an Année Recherche from AP-HP and by a SNFMI fellowship.

## Author contributions

Conceptualization: P.C., A.S., S.F., JC.W., CA.R., and M.M.; Data curation: P.C., A.S; Formal Analysis: A.S., P.C., A.R., I.A., A.V., M.B., S.H., A.B., F.R., G.B.; Funding acquisition: S.F., JC.W., CA.R., and M.M.; Investigation: A.S., P.C., A.R., I.A., A.V.; Methodology: A.S., JC.W., CA.R., P.C. and M.M.; Project administration: P.C., S.B., S.F., JC.W, CA.R., and M.M.; Resources: JM.P., F.R., S.F., E.C., CA.R., M.M.; Software: P.C; Supervision: P.C., JC.W., CA.R, and M.M.; Validation: A.R, I.A and A.V.; Visualization: P.C., A.S., I.A., A.R., A.V., M.M.; Writing – original draft: P.C., A.S., JC.W, CA.R., and M.M.; Writing – review & editing: all authors.

## Declaration of interests

M.M. received research funds from GSK, outside of the submitted work and personal fees from LFB and Amgen, outside of the submitted work. JC.W. received consulting fees from Mérieux, outside of the submitted work. JM.P. received personal fees from Abbvie, Gilead, Merck, and Siemens Healthcare, outside the submitted work.

## Methods

| Reagent or resource | Source | Identifier |
| --- | --- | --- |
| **Antibody** |  |  |
| CD3 | Biolegend | UCHT1 ; RRID : AB_830754 |
| CD14 | BD Bioscience | M φP9 ; RRID : AB_1645464 |
| CD19 | BD Bioscience | HIB19 ; RRID : AB_11154408 |
| CD38 | BD Bioscience | HIT2 ; RRID AB_394184 |
| CD27 | Biolegend | M-T271 ; AB_2562598 |
| CD11c | BD Bioscience | S-HCL-3 ; AB_2742232 |
| IgD | Life technologies | Polyclonal, RRID : AB_2536563 |
| His Tag | Biolegend | J095G46 ; RRID:AB_2563634 |
| His Tag | Biolegend | J095G46 ; RRID:AB_2715818 |
| CD71 | Biolegend | CY1G4 ; RRID:AB_2563118 |
| CD21 | BD Bioscience | B-ly4; RRID:AB_2741028 |
| CD14 | BD Bioscience | M5E2 ; RRID:AB_396590 |
| CD15 | BD Bioscience | HI98 ; RRID:AB_395802 |
| CD56 | BD Bioscience | B159 ; RRID:AB_395906 |
| IgD | BD Bioscience | IA6-2 ; RRID:AB_396114 |
| CD3 | BD Bioscience | UCHT1 ; RRID:AB_10561682 |
| T-Bet | Biolegend | 4B10 ; RRID:AB_2561760 |
| CD38 | Biolegend | HIT2 ; RRID:AB_314361 |
| CD38 (Totalseq-C) | Biolegend | HB-7; RRID:AB_2820007 |
| CD138 (Totalseq-C) | Biolegend | MI15; RRID:AB_2810567 |
| CD27 (Totalseq-C) | Biolegend | O323; RRID:AB_2800747 |
| CD11c (Totalseq-C) | Biolegend | S-HCL-3; RRID:AB_2801018 |
| CD21 (Totalseq-C) | Biolegend | Bu32; RRID:AB_2800953 |
| CD71 (Totalseq-C) | Biolegend | CY1G4; RRID:AB_2800885 |
| CXCR4 (Totalseq-C) | Biolegend | 12G5; RRID:AB_2800791 |
| CXCR3 (Totalseq-C) | Biolegend | G025H7; RRID:AB_2800949 |
| CD69 (Totalseq-C) | Biolegend | FN50; RRID:AB_2800810 |
| HLA-DR (Totalseq-C) | Biolegend | L243; RRID:AB_2800795 |
| CD307d (FCRL4; Totalseq-C) | Biolegend | 413D12; RRID:AB_2832666 |
| CD307e (FCRL5; Totalseq-C) | Biolegend | 509f6; RRID:AB_2819969 |
| CD95 (Fas; Totalseq-C) | Biolegend | DX2; RRID:AB_2800787 |
| Integrin B7 | Biolegend | FIB504; RRID:AB_2810481 |
| CD49d (Integrin alpha 4; Totalseq-C) | Biolegend | 9F10; RRID:AB_2814137 |
| CD103 (Totalseq-C) | Biolegend | Ber-ACT8; AB_2800933 |
| LAG-3 (Totalseq-C) | Biolegend | 11C3C65; RRID:AB_2814327 |
| TIM3 (Totalseq-C) | Biolegend | F38-2E2; RRID:AB_2800925 |
| CD305 (LAIR1; Totalseq-C) | Biolegend | NKTA255; RRID:AB_2832668 |
| NKG2D (Totalseq-C) | Biolegend | 1D11; RRID:AB_2800844 |
| CD73 (Totalseq-C) | Biolegend | AD2; RRID:AB_2800916 |
| His Tag (Totalseq-C) | Biolegend | Custom conjugation |
| His Tag (Totalseq-C) | Biolegend | Custom conjugation |
| **Biological samples** |  |  |
| Cryopreserved PBMCs from S-Cov and M-Cov patients | Henri Mondor Hospital, Assistance Publique des Hôpitaux de Paris |  |
| Cryopreserved PBMCs from pre-pandemic subjects | French Blood Agency (EFS) |  |
| **Chemical, peptides, and recombinant proteins** |  |  |
| Sars-CoV-2 Spike | Pasteur Institute |  |
| Sars-CoV-2 RBD | Pasteur Institute |  |
| HKU1 Spike | Pasteur Institute |  |
| OC43 Spike | Pasteur Institute |  |
| Live dead aqua | Life technologies | L34957 |
| Recombinant human IL-2 | PeproTech | 200-02 |
| Recombinant human IL-4 | PeproTech | 200-04 |
| Recombinant human IL-21 | PeproTech | 210-21 |
| Recombinant human BAFF | PeproTech | 310-13 |
| Oligonucleotide Conjugation Kit | Abcam | ab218260 |
| **Softwares and algorithms** |  |  |
| Kaluza v2.1 | Beckman Coulter | https://www.beckman.fr |
| Flowjo v10.7.1 | FlowJo, LLC | https://www.flowjo.com |
| GraphPad Prism v8 | GraphPad | https://www.graphpad.com |
| Codon Code Aligner v9 | Codon Code Corporation | https://www.codoncode.com/ |
| Python v3.7.6 | Python Software Foundation | https://www.python.org/ |
| R v4.0.2 | R Foundation | https://www.r-project.org |
| RStudio v1.3.1056 | RStudio | https://rstudio.com |
| IgBLASTn v1.16.0 | NCBI | https://www.ncbi.nlm.nih.gov/igblast/ |
| Docker desktop v2.5.0.0 | Docker, Inc | https://www.docker.com/products/docker-desktop |

## RESOURCE AVAILABILITY

### Lead Contact

Further information and requests for resources and reagents should be directed to and will be fulfilled by the Lead Contact, Matthieu Mahévas.

### Materials Availability

No unique materials were generated for this study.

### Data and Code Availability

All NGS data reported in this study will be made available.

## EXPERIMENTAL MODEL AND SUBJECT DETAILS

### Study participants

In total 50 patients with COVID-19 were recruited. We included 21 COVID-19 patients requiring oxygen (S-CoV) and 18 healthcare workers, with a mild COVID-19 (M-CoV). SARS-CoV-2 infection was defined as confirmed reverse transcriptase polymerase chain reaction (RT-PCR) on nasal swab or clinical presentation associated with typical aspect on CT-scan and/or serological evidence. Samples were collected in mean 18.8 (±SD: 8.8 days) days after disease onset in S-CoV, and in mean 35.5 days (± SD: 12.8 days) for M-CoV. Samples were additionally collected 3 months (mean ±SD: 89.8 ± 15.8 days for S-CoV and 94.2 ± 8.2 for M-CoV), and 6 months (mean ± SD: 170.5 ± 12.9 days for S-CoV and 184.6 ± 10.2 for M-CoV) after onset. Clinical and biological characteristics of these patients are summarized in Table S1. Patients were recruited at Henri Mondor University Hospital (AP-HP), between March and May 2020. Samples from healthy donors were obtained at EFS Henri-Mondor and frozen before 2019. MEMO-COV-2 study (NCT04402892) was approved by the ethical committee Ile-de-France VI (Number: 40-20 HPS), and was performed in accordance with the French law. Written informed consent was obtained for all participants.

## METHOD DETAILS

### SARS-CoV-2 viral detection by RT-PCR

Nasopharyngeal (NP) swabs in transport media and plasma samples were held at 4 °C (or −80°C if > 12h) prior to testing on the Alinity M SARS CoV-2 Amp kit and/or Cepheid Xpert Xpress SARS-CoV-2 kit. The GeneXpert ®Dx System (Cepheid) and Alinity M (Abbott) perform automated specimen processing and real-time RT-PCR analysis. The Cepheid Xpert Xpress SARS-CoV-2 kit detects N and E genes of the SARS CoV-2 genome; Alinity M SARS CoV-2 Amp kit detects RdRp and N genes.

### Anti-S and anti-N commercial assays

Serological assays were performed for IgG anti-N, IgG anti-S and total antibodies (ab) anti-S detection. Serum samples were processed for anti-Nucloprotein (N) detection on Abbott SARS-CoV-2 IgG chemiluminescent microparticle immunoassay following the manufacturer’s instructions. Serum samples were further analyzed using the VITROS Immunodiagnostic Products, for IgG Anti-Spike (S) SARS-CoV-2 detection and for total Ab anti-spike (S) detection (Ortho Clinical Diagnostics). All assays were performed by trained laboratory technicians according to the respective manufacturer's standard procedures. Qualitative results and index values reported by the instruments were used in analysis.

### Recombinant proteins purification

#### Construct design

The human coronavirus Spike (S) proteins were expressed as ectodomains that were stabilized to preserve their trimeric prefusion conformation. The SARS-CoV-2 S (residues 1-1208) was stabilized by introducing six proline substitutions (F817P, A892P, A899P, A942P, K986P, V987P), a GSAS substitution at the furin cleavage site (residues 682-685) and a C-terminal Foldon trimerization motif (Hsieh et al., 2020), followed by Hisx8 and Strep tags. This construct was cloned using its endogenous signal peptide in pcDNA3.1(+). The OC43 S ectodomain (residues 15-1263) was cloned in the pCAGGS vector with a CD5 N-terminal signal peptide and a C-terminal GCN4 trimerization motif, a thrombin cleavage site and a single Strep-tag. Additionally, it contains mutations to abolish the furin cleavage (754-RRSRG-758 to 754-GGSGG-758) at the S1-S2 junction (Tortorici et al., 2019). The HKU1-CoV S protein (residues 14-1276) was stabilized by a double-proline mutation (N1067P, L1068P), the substitution of the RRSRG motif by GGSGG (residues 754-758) to avoid a potential S1/S2 furin cleavage, and a C-terminal Foldon motif. This construct was cloned in pcDNA3.1(+) with an IgK signal peptide and a thrombin cleavage site at the C-terminus that was followed by a Hisx8 tag.

The SARS-CoV-2 Receptor Binding Domain (RBD) was cloned in pcDNA3.1(+) encompassing residues 331-528 from the Spike ectodomain, and it was flanked by an N-terminal IgK signal peptide and a C-terminal Thrombin cleavage site followed by a Hisx8-tag.

#### Protein expression and purification

The plasmids coding for the recombinant proteins were transiently transfected in Expi293F™ cells (Thermo Fischer) using FectroPRO® DNA transfection reagent (Polyplus), according to the manufacturer’s instructions. The cells were incubated at 37 °C for 5 days and then the culture was centrifuged and the supernatant was concentrated. The proteins were purified from the supernatant by affinity and size-exclusion chromatography (SEC). The first purification step was performed using StrepTactin columns (IBA) (SARS-Cov-2 S, OC43 S) or His-Trap™ Excel columns (GE Healthcare) (HKU1 S, SARS-Cov-2 RBD). The different S ectodomains were further purified using a Superose6 10/300 colum (GE Healthcare) equilibrated in PBS, while a Superdex200 10/300 column was used for the SARS-Cov-2 RBD.

### Flow cytometry and cell sorting

PBMCs were isolated from venous blood samples via standard density gradient centrifugation and used after cryopreservation at −150°C. Cells were thawed using RPMI-1640 (Gibco) 10% FBS, washed twice and incubated with 10μg of the SARS-CoV-2 his tagged spike protein in 100μL of PBS (Gibco) 2% FBS during 20 minutes on ice. Cells were washed and resuspended in the same conditions, then fluorochrome-conjugated antibody cocktail including the 2 anti-His was added at pre-titrated concentrations for 20 min at 4°C and viable cells were identified using a LIVE/DEAD Fixable Aqua Dead Cell Stain Kit (Thermo Fisher Scientific) incubated with conjugated antibodies (see Table S3). If a permeabilization was needed (T-Bet staining), cells were washed and resuspended in 250μL of Fix/perm (eBioscience) for 25 minutes, then washed with appropriate buffer before adding the conjugated antibody on the cell pellet for a further 25 minutes at 4°C. Samples were acquired using a LSR Fortessa SORP (BD Biosciences). For cell sorting, cells were stained using the same protocol then sorted in 96 plates using an ultra-purity mode on a MA900 cell sorter (SONY), or Aria III (BD Biosciences). Data were analyzed with FlowJo or Kaluza softwares. Detailed gating strategies for individual markers are depicted in Figure S1.

For UMAP generation and visualization (Figure 3A-C), data from all 83 samples from patient with complete panel acquisition at M0, M3 and M6 (Table S1) in our dataset were individually down-sampled to 3000 cells each using the Downsample (v3.3) plugin in FlowJO. All samples were subsequently concatenated and FlowJO’s UMAP (v3.1) plugin was used to calculate the UMAP coordinates for the resulting 249,000 cells (with 30 neighbors, metric = euclidian and minimum distance = 0.5 as default parameters). FlowSOM plugin was used in parallel on the same downsampled dataset to create a self-organizing map (using n = 9 clusters as default parameter) that was then applied to the initial FCS files from all 83 samples to calculate total and spike-specific memory B cell repartition in identified clusters. Both UMAP and FlowSOM plugin were runned taking into account fluorescent intensities from the following parameters: FSC-A, SSC-A, CD19, CD21, CD11c, CD71, CD38, CD27 and IgD. Contour plots (equal probability contouring, log scale) for each identified cluster representing more than 5% of the total cells in the dataset were further overlaid in FlowJO for visualization purposes.

### Single-cell culture

Single cell culture was performed as previously described (Crickx et al., 2019). Single B cells were sorted in 96-well plates containing MS40 cells expressing CD40L (kind gift from G. Kelsoe). Cells were cocultured at 37°C with 5% CO2 during 21 or 25 days in RPMI-1640 (Invitrogen) supplemented with 10% HyClone FBS (Thermo Scientific), 55 μM 2-mercaptoethanol, 10 mM HEPES, 1 mM sodium pyruvate, 100 units/mL penicillin, 100 ug/mL streptomycin, and MEM non-essential amino acids (all Invitrogen), with the addition of recombinant human BAFF (10 ng/ml), IL2 (50 ng/ml), IL4 (10 ng/ml), and IL21 (10 ng/ml; all Peprotech). Part of the supernatant was carefully removed at days 4, 8,12, 15 and 18 and the same amount of fresh medium with cytokines was added to the cultures. After 21 days of single cell culture, supernatants were harvested and stored at −20°C. Cell pellets were placed on ice and gently washed with PBS (Gibco) before being resuspended in 50μL of RLT buffer (Qiagen) supplemented with 10% beta-mercaptoethanol and subsequently stored at −80°C until further processing.

### ELISA

IgG and SARS-CoV 2, HKU1-CoV and OC43-CoV spike-specific IgG from culture supernatants were detected by home-made ELISA. Briefly, 96 well ELISA plates (Thermo Fisher) were coated with either goat anti-human Ig (10μg/ml, Invitrogen) or recombinant SARS-CoV-2, HKU1 or OC43 spike or SARS-CoV-2 RBD protein (2.5μg/ml each) in sodium carbonate during 1h at 37°C. After plate blocking, cell culture supernatants were added for 1h, then ELISA were developed using HRP-goat antihuman IgG (1μg/ml, Immunotech) and TMB substrate (Eurobio). OD450 and OD620 were measured and Ab-reactivity was calculated after subtraction of blank wells. Supernatants whose ratio of OD450-OD620 over control (i.e. non spike specific supernatant from the same single cell culture assay) was over 3 were considered as positive for RBD ELISA. A cut-off at a ratio of 5 was chosen for HKU1 and OC43 ELISA.

### Single-cell IgH sequencing

Clones whose culture had proved successful (IgG concentration ≥ 1μg/mL at day 21-25) were selected and extracted using the NucleoSpin96 RNA extraction kit (Macherey-Nagel) according to the manufacturer’s instruction. A reverse transcription step was then performed using the SuperScript IV enzyme (ThermoFisher) in a 14 μl final volume (42°C 10min, 25°C 10min, 50°C 60min, 94°C 5min) with 4 μl of RNA and random hexamers (GE Healthcare). A PCR was further performed based on the protocol established by Tiller et al (Tiller et al., 2008). Briefly, 3.5 μl of cDNA was used as template and amplified in a total volume of 40 μl with a mix of forward L-VH primers (Table S3) and reverse Cγ primer and using the HotStar® Taq DNA polymerase (Qiagen) and 50 cycles of PCR (94°C 30s, 58°C 30s, 72°C 60s). PCR products were sequenced with the reverse primer CHG-D1 and read on ABI PRISM 3130XL genetic analyzer (Applied Biosystems). Sequence quality was verified with the CodonCode Aligner software (CodonCode Corporation) and data were analyzed with the IMGT/HighV-QUEST web portal (from The International Immunogenetics Information System) or in parallel with the VDJ sequences generated as part of our scRNA-seq dataset (see below).

### Oligonucleotide conjugation of anti-His antibody

Unconjugated anti-His antibodies were purchased at Biolegend and custom oligonucleotides were ordered at Integrated DNA Technologies, following the 10Xgenomics protocol available at https://support.10xgenomics.com/single-cell-gene-expression/overview/doc/demonstratedprotocol-cell-surface-protein-labeling-for-single-cell-rna-sequencing-protocols and the barcode whitelist. Sequences of the 2 HPLC purified barcoded oligonucleotides were:/5AmMC12/CGGAGATGTGTATAAGAGACAGNNNNNNNNNNTGCATAGCCTGTGGANNNNNNNNNCCCATATAAGAAA and/5AmMC12/CGGAGATGTGTATAAGAGACAGNNNNNNNNNNCTCTCCAATGTACTCNNNNNNNNNCCCATATAAGAAA. Oligonucleotide antibody conjugation was performed using the Oligonucleotide Conjugation Kit from Abcam, according to the manufacturer instructions using an oligonucleotide/antibody ratio of 5:1.

### Single-cell RNA-seq library preparation and sequencing

Peripheral (CD3^−^CD14^−^CD15^−^CD56^−^CD19^+^IgD^−^) B cells were FACS-sorted (MA900, Sony) in PBS/0.08% FCS from 4 patients (S-CoV) at baseline (M0) and 6 months (M6). 5×10^4^ to 10×10^5^ cells were obtained for each subset. The scRNA-seq libraries were generated using Chromium Next GEM Single Cell V(D)J Reagent Kit v.1.1 with Feature Barcoding (10x Genomics) according to the manufacturer’s protocol. Gene expression (mRNA), ADT and VDJ BCR libraries were constructed. Briefly, cells were counted and up to 20 000 cells were loaded in the 10x Chromium Controller to generate single-cell gel-beads in emulsion. After reverse transcription, gel-beads in emulsion were disrupted. Barcoded complementary DNA was isolated and amplified by PCR. Following fragmentation, end repair and A-tailing, sample indexes were added during index PCR. The purified libraries were sequenced on a Novaseq S2 flowcell (Illumina) with 26 cycles of read 1, 8 cycles of i7 index and 91 cycles of read 2, targeting a median depth of 50000 reads per cell for gene expression and 5000 reads per cell for each other two libraries (BCR VDJ and ADT Feature barcoding).

### Single-cell gene expression analysis

Paired-end FASTQ reads for all three libraries were demultiplexed and aligned against the GRCh38 human reference genome (GENCODE v32/Ensembl 98; July 2020) using 10x Genomics’ Cell Ranger v4.0.0 pipeline. Outputs of Cell Ranger were directly loaded into Seurat v3.2.2 (Stuart et al., 2019) for further QC steps and analysis. Following manual inspection of cell quality, only genes detected in at least 10 cells and cells with more than 500 unique genes detected and less than 25% of UMI counts mapped to mitochondrial genes were kept. Reads mapping to the immunoglobulin genes locus were further stored in a separate assay at this step to avoid unwanted clustering based solely on differential isotype expression and cells with exactly one heavy chain sequence were retained for final analysis. UMI counts were then log-normalized. The top 2,000 highly variable genes were identified using the FindVariableFeatures() function in Seurat and the default vst method. Normalized counts were then scale end centered using the ScaleData() function, removing unwanted variation related to the percentage of mitochondrial UMI counts at that step. After principal component analysis, potential donor and sort-specific batch effects were removed using the Harmony algorithm (Korsunsky et al., 2019). The first 30 corrected PCA dimensions were then used to construct a knn graph (k=20 neighbors) and perform graph-based clustering (Louvain) with a resolution parameter of 0.2 as well as compute the UMAP coordinates for each cell. Further separation of the “Activated” cell cluster was performed using a resolution parameter of 0.4. Two small clusters with T (0.2%) and monocyte (0.1%) signatures were removed from further analysis as clear doublets, as well as a separate cluster of apoptotic cells (2.1% of all cells), grouped solely based on high percentages of mitochondrial UMI count.

### Computational analyses of VDJ sequences

Processed FASTA sequences from cultured single-cell heavy chain sequencing, 10x single-cell RNA sequencing and 874 published SARS-CoV-2 RBD and/or S-specific antibodies (Brouwer et al., 2020; Kreer et al., 2020; Liu et al., 2020; Robbiani et al., 2020; Seydoux et al., 2020; Shi et al., 2020; Wec et al., 2020; Zost et al., 2020) were annotated using Igblast v1.16.0 against the human IMGT reference database. Non-productively rearranged sequences were removed at that step as well as sequences from cells that did not pass the initial QC cut-offs from our scRNA-seq analysis. Cases of 10x barcodes with two or more consensus heavy chain sequences for which more than ten UMI were detected were generally flagged as potential doublets for removal from our scRNA-seq analysis. Similarly, cases where no clear heavy chains could be attributed (none above 10 UMIs) were also flagged for removal. Two exceptions were made: 1/ in cases of identical CDR3s but differing isotypes (c_call), in which case the isotype switched sequence was kept and UMI counts from both contigs were aggregated; and 2/ in cases when one the heavy chains was clearly over represented at the UMI level and the second most represented sequences did not exceed ten UMIs, in which case the most represented sequence was kept.

Clonal cluster assignment (DefineClones.py) and germline reconstruction (CreateGermlines.py) was performed using the Immcantation/Change-O toolkit (Gupta et al., 2015) on all heavy chain V sequences. Sequences that had the same V-gene, same J-gene, including ambiguous assignments, and same CDR3 length with maximal length normalized nucleotide hamming distance of 0.15 were considered as potentially belonging to the same clonal group. Mutation frequencies in V genes were then calculated using the calcObservedMutations() function from Immcantation/SHazaM v1.0.2 R package. For the analysis of the initial ASC response in our 10x dataset (**Figure 2E/F**), clonal assignments were further corrected using available light chain information (light_cluster.py script from Immcantation). All SARS-CoV-2 S-specific clones described in **Figure 4** were manually curated based on available light chain information (10x and published antibodies only) and CDR3 sequences. Further clonal analysis were implemented in R. Based on final clonal affectation, clones were defined as SARS-CoV-2 S specific if they contained 1 or more validated single-cell culture sequence or if more than ten percent of the cells from that clone were positively stained by our barcoded His-tagged S protein in our scRNAseq dataset. Clones were defined as SARS-CoV-2 RBD or HKU1/OC43 S-specific if they contained 1 or more validated single-cell culture sequence with a positive ELISA against one of these proteins. Graphics were obtained using the ggplot2 v3.3.2 and circlize v0.4.10 packages. Phylogenetic trees were generated using the Immcantation/IgPhyML toolkit (Immcantation/suite v4.0.0 docker image) and further visualize in R using the Alakazam v1.0.2 and igraph v1.2.6 packages.

### Virus neutralization assay

Virus neutralization was evaluated by a focus reduction neutralization test (FRNT). Vero E6 cells were seeded at 2×10^4^cells/well in a 96-well plate 24h before the assay. Two hundred focus-forming units (ffu) of Sars-CoV-2 virus (BetaCoV/France/IDF0372/2020 strain, a kind gift from the National Reference Centre for Respiratory Viruses at Institut Pasteur, Paris, originally supplied through the European Virus Archive goes Global platform) were pre-incubated with serial dilutions of heat-inactivated sera or supernatant from B-cell clones for 1h at 37°C before infection of cells. After 2h infection, the virus/antibody mix was removed and foci were left to develop in presence of 1.5% methylcellulose for 2 days. Cells were then fixed with 4% formaldehyde and foci were revealed using an anti-S antibody and a secondary HRP-conjugated secondary antibody. Foci were visualized by diaminobenzidine (DAB) staining and counted using an Immunospot S6 Analyser (Cellular Technology Limited CTL). Neutralization curves and 50% FRNT values were calculated by nonlinear regression analysis using Prism 6, GraphPad software.

### Statistics

Kruskal-Wallis test and Mann-Whitney test were used to compare continuous variables as appropriate. A *P*-value ≤ 0.05 was considered statistically significant. Statistical analyses involved use of GraphPad Prism 8.0 (La Jolla, CA, USA).

**Figure S1.**
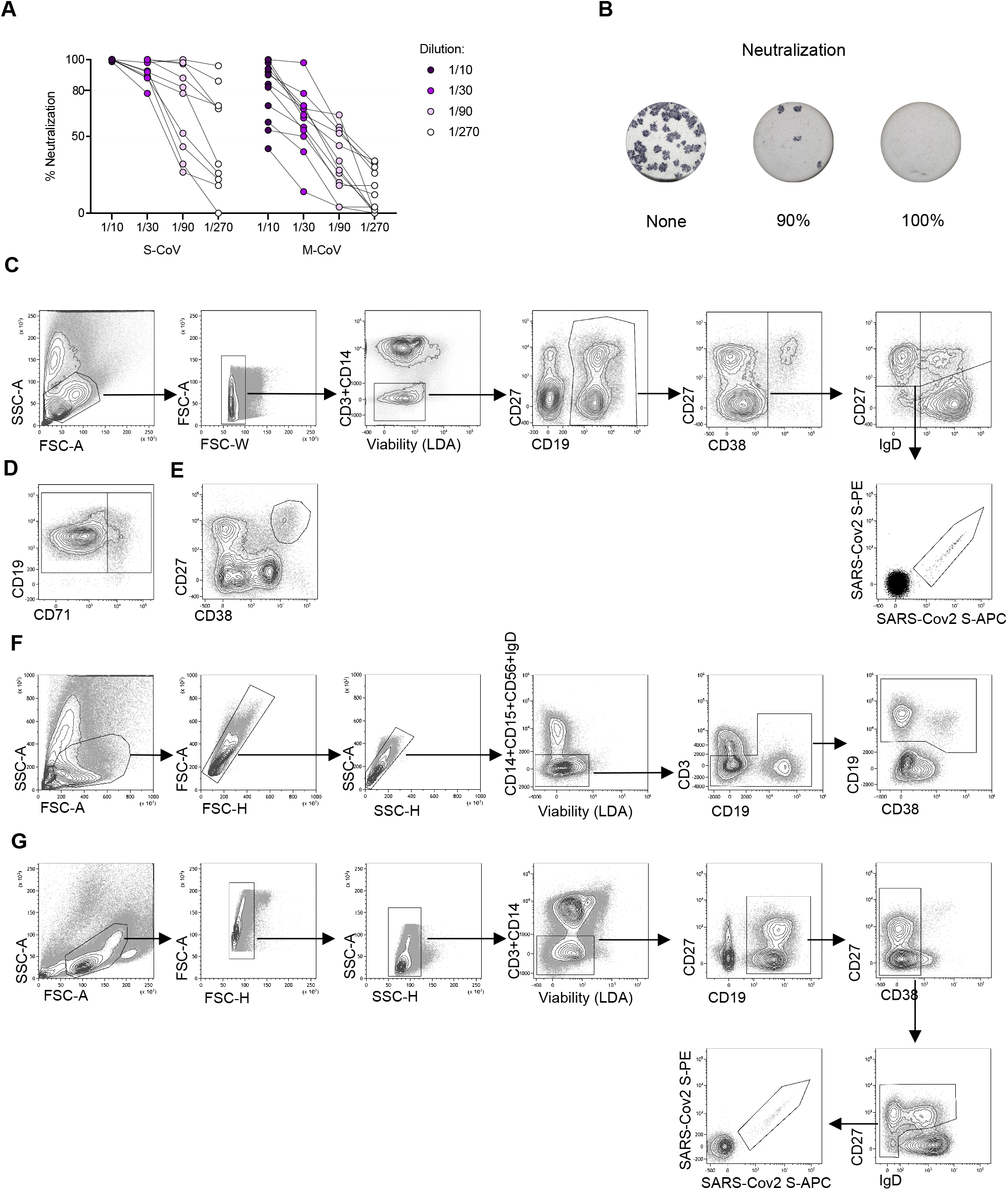
Quantification and functional assessment of the anti-SARS-CoV2 humoral immune response in COVID-19 convalescent patients (Related to Figure 1). **(A)** Percentage of in vitro neutralization shown by individual sera from S-CoV (n=10) and M-CoV (n=12) at M6 at increasing dilutions. **(B)** Representative wells for the neutralization assay. Blue spots represent SARS-CoV-2 positive cells. **(C-G)** Flow cytometric gating strategies for the analysis and sorting of major B cell population from PBMCs of convalescent COVID-19 patients. **(C)** Gating strategy to analyze SARS-CoV2 S-specific B cell population. Lymphocytes were first gated based on morphology, before exclusion of doublets, dead cells and CD3/CD14 cells. CD19^+^ cells were next gated before exclusion of CD38^hi^ plasma cells. CD38^int/-^ cells were then divided in four quadrants using CD27 and IgD. Upper left quadrant defines memory B cells (MBCs), lower left quadrant double-negative (DN), upper right quadrant CD27^+^IgD^+^ cells (MZB) and lower right quadrant naive B cells (excluding CD38^hi^ transitional). SARS-CoV2 S-specific B cells were then analyzed within the B cell population of interest using a His-tagged SARS-CoV2 S double staining strategy. **(D)** Gating strategy to separate activated and classical switched B cells using CD71, within the IgD^−^CD27^+^ gate **(E)** Gating strategy to analyze CD27^hi^CD38^hi^ plasma cells in CD3^−^CD14^−^ live cells. **(F)** Gating strategy for sorting of CD19^+^IgD^−^ cells from PBMCs for 10X-Chromium single cell experiment. Lymphocytes were gated before exclusion of doublets and of CD14/CD15/CD56/IgD^+^ cells, before sorting of CD19^+^ cells. **(G)** Gating strategy for single-cell sorting of SARS-CoV2 S-specific B cells for single cell culture at M3 and M6. Lymphocytes were gated before double exclusion of doublets, dead cells and CD3/CD14 cells. CD19^+^ cells were then gated before exclusion of CD38^hi^ plasma cells. CD38^int/-^ cells were further divided into four quadrants using CD27 and IgD. Double stained SARS-CoV2 S specific cells were sorted from all three non-naïve quadrants (not IgD^+^CD27^−^).

**Figure S2.**
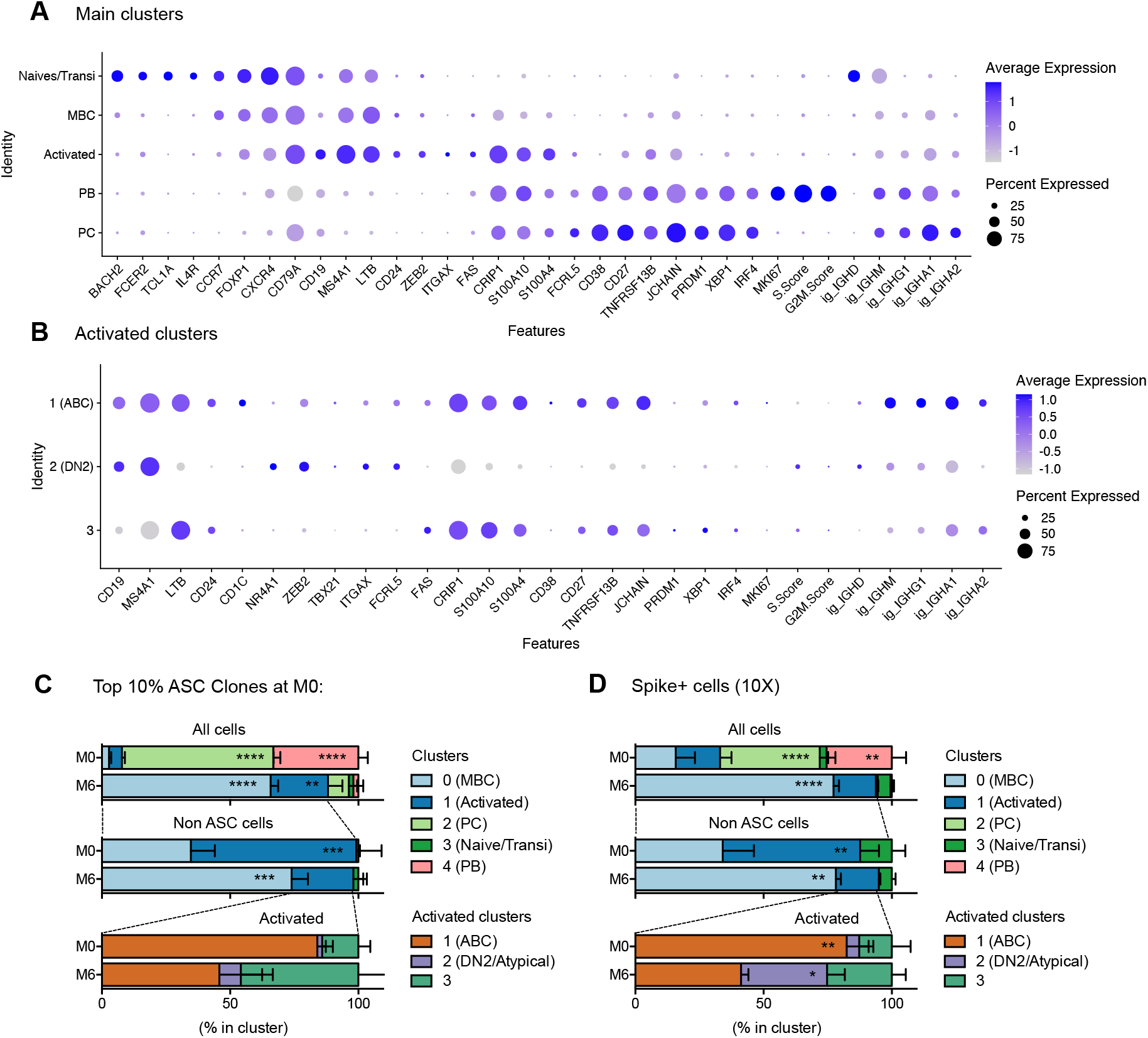
Gene expression markers and cluster assignments for the main single cell populations identified in acute and convalescent S-COV patients (Related to Figure 2). **(A-B)** Dot plots showing expression of selected genes in cells from the main clusters **(A)** or in cells from the “Activated” cluster **(B)**. Size of dots represents the percentage of cells in the cluster in which transcripts for that gene are detected. Dot color represents the average expression level (scaled normalized counts) of that gene in the population. **(C-D)** Relative cluster distribution at M0 and M6 for cells belonging to one of the 10 percent most expanded antibody secreting cell (ASC) clones **(C)** or positive for barcoded-SARS-CoV2-S staining **(D)**. Top panels represent cluster distribution for all cells, middle panels represent cluster distribution for non-ASC cells and bottom panels represent cluster distribution for cells belonging to the “Activated” cluster. Bars indicate mean with SEM.

**Figure S3.**
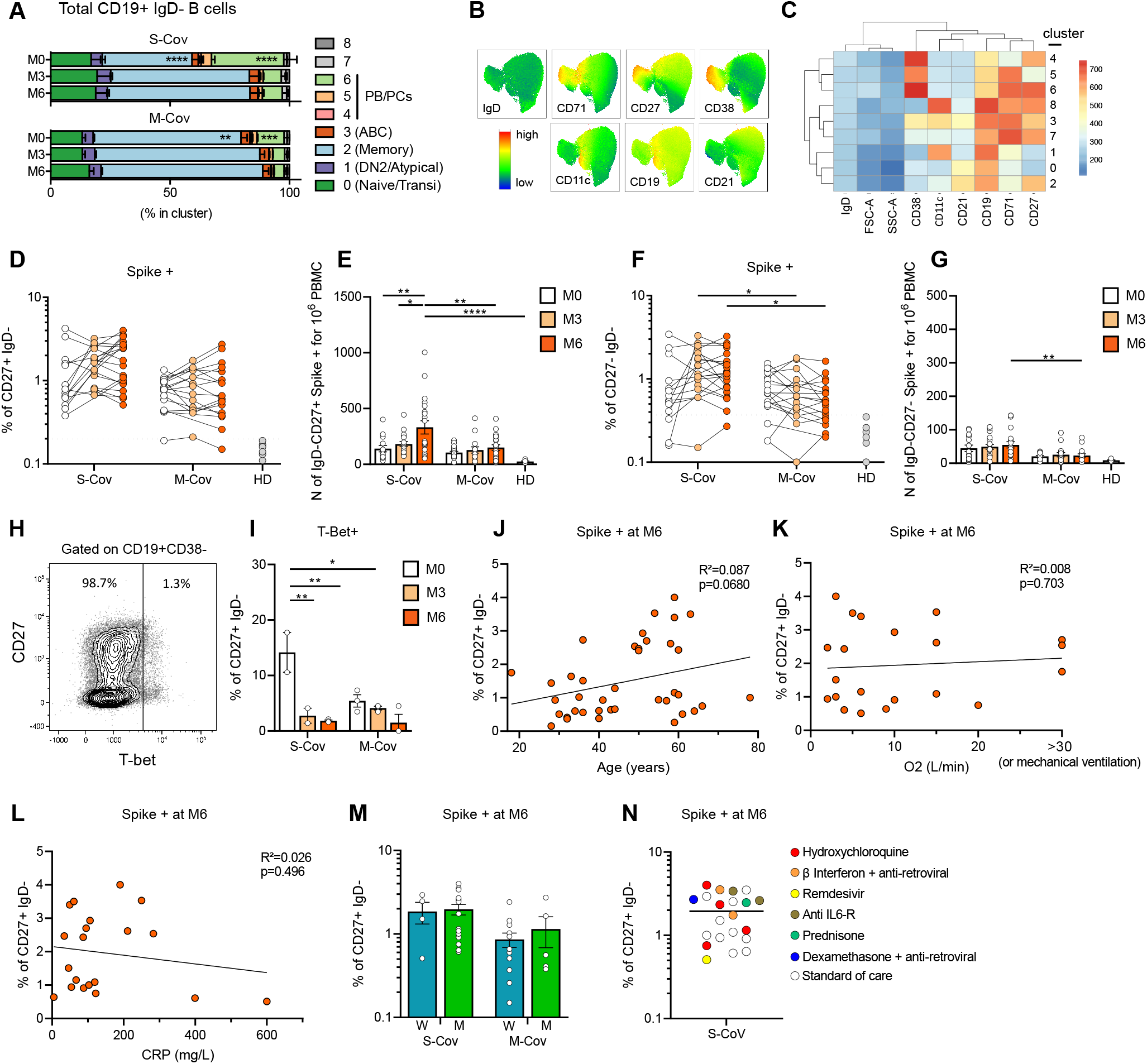
Phenotypic characterization of total and SARS-CoV2 S-specific B cell populations in convalescent COVID-19 patients (Related to Figure 3). **(A)** Cluster distribution of IgD^−^ CD19^+^ B cell across FlowSOM identified clusters in concatenated FCS files from S-CoV (n=15) and 16 M-CoV (n=16) donors (Table S1) and all time points (M0, M3 and M6) of our cohort. **(B)** UMAP overlaid scaled intensity for IgD, CD71, CD27, CD38, CD11c, CD19, CD21 expression in each cell from the compiled FCS. **(C)** Heat map representing the mean fluorescent intensity for IgD, CD71, CD27, CD38, CD11c, CD19, CD21 in each identified FlowSOM cluster. **(D-E)** Percentage (D) and absolute numbers (E) of SARS-CoV2 S-specific CD27^+^IgD^−^ MBCs at indicated time points in S-CoV and M-CoV patients. **(F-G)** Percentage (F) and absolute number (G) of SARS-CoV2 S-specific cells among CD27^−^IgD^−^ DN cells. (D and F) Each dot is a patient and lines show the corresponding dot at other time points. Dashed line indicates the threshold based on healthy donors (mean + 2 SD). (E and G) Bars indicate the mean with SEM. **(H)** Representative dot plot for T-bet staining in CD19^+^CD38^−^ B cells from COVID-19 patients. **(I)** Percentage of intra cellular T-Bet staining in SARS-CoV2 S-specific CD19^+^IgD^−^CD27^+^CD38^−^ cells from S-CoV (n=2) and M-CoV (n=3) patients at indicated time points. Bar indicate the mean with SEM. **(J, K, L)** Correlation between the number of SARS-CoV2 S-specific CD27^+^IgD^−^ MBCs at M6 and age at diagnosis (J), maximum oxygen flow during hospitalization (liters/minutes, only in S-CoV) (K) and maximum CRP during hospitalization (mg/L, available only in S-CoV) (L) using linear regression. **(M)** Number of spike-specific CD27^+^IgD^−^ MBCs at M6 according to the initial disease severity (S-CoV and M-CoV) and the sex of the patient (Women, W, blue and Men, M, blue). Bars indicate the mean with SEM. **(N)** Plot showing the number of spike specific CD27^+^IgD^−^ cells in all the n=21 S-CoV patients analyzed at M6 colored according to the treatment received during initial hospitalization. Standard of care included oxygen, low dose anticoagulant, antibiotic if needed and symptomatic treatments. Anova and two-tailed Mann-Whitney tests were performed (**P < 0.01, *P < 0.05).

**Figure S4.**
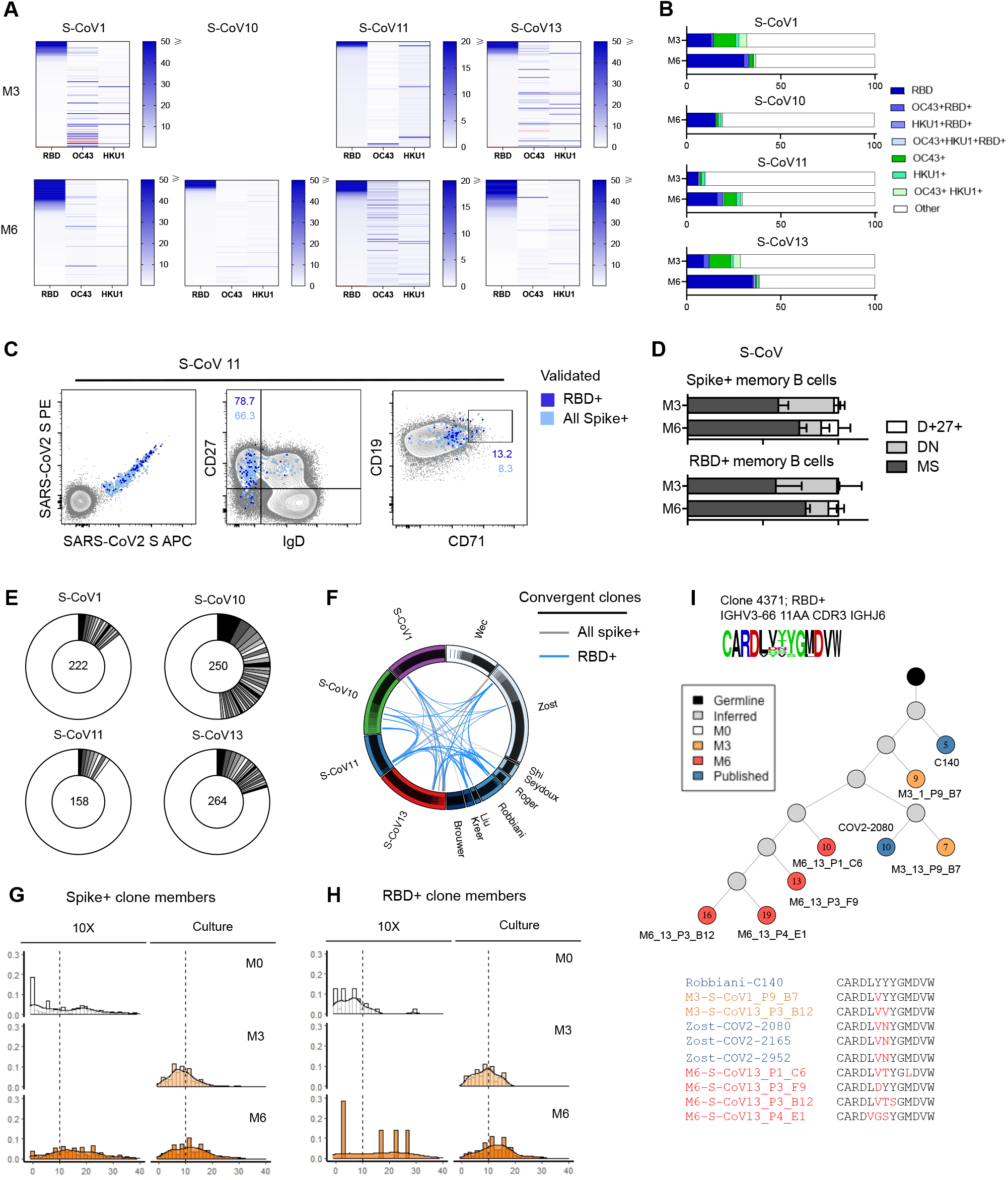
Cross-reactivity, convergence and accumulation of somatic mutations in anti-SARS-CoV2 S-specific memory B cells from convalescent COVID-19 patients (Related to Figure 4). **(A)** Heat map showing the RBD, OC43 and HKU1 ELISA blank ratio for all tested SARS-CoV2-specific single cell culture supernatant at M3 and M6. Each line represents one tested supernatant. Light red lines indicate that no value is available for that cell **(B)** Bar plot showing the proportion of RBD specificity and cross-reactive specificities (RBD^+^OC43^+^, RBD^+^HKU1^+^OC4^+^, RBD^+^HKU1^+^, OC43^+^, HKU1^+^, OC43^+^HKU1^+^) among S-specific cells for each patient. **(C)** FACS plot representing index sorting data of spike-specific cells according to their specificity for RBD. **(D)** Repartition of the spike- and RBD-specific cells at M3 and M6 between the MBCs, DN or IgD^+^CD27^+^ compartments. **(E)** Pie chart representing clone size in all the sequences generated via single cell culture for each of the 4 patients previously included in our scRNAseq analysis. Total number of sequenced cells is indicated in the middle of the pie. **(F)** Circus plot showing SARS-CoV2 S-(grey) or RBD-(light blue) specific clones shared between patients in our dataset or containing sequences from the literature. **(G-H)** Histograms showing the distribution of mutations in Ig VH for SARS-CoV2 S-(G) and RBD-(H) specific clones at M0, M3 and M6 according to the assay of origin. **(I)** Evolutionary tree of a convergent RBD-specific and neutralizing clone, built on sequences from 10X scRNA-seq and cell culture from two patients and from the literature. Each circle represents a unique sequence from that clone. Circle color indicates time-point of origin and the number inside indicates the calculated number of mutations from inferred germline. Grey indicates a theoretically inferred common progenitor. *indicate that the antibody associated with that sequence has been validated as neutralizing in vitro. CDR3 from all sequences in the tree are represented as a frequency plot logo (Top left) as well as below the tree, where each amino-acid in red indicates a change compared to the first listed CDR3 sequence.

